# Genome-wide mapping of histone modification H3K4me3 in bovine oocytes and early embryos

**DOI:** 10.1101/2021.11.22.469629

**Authors:** Yanna Dang, Lei Luo, Yan Shi, Shuang Li, Shaohua Wang, Kun Zhang

**Author notes:** To whom correspondence may be addressed., **Corresponding Author:** Kun Zhang, Room 301 E Building, 866 Yuhangtang Rd, Hangzhou, Zhejiang 310058, China.

## Abstract

Reprogramming of histone modifications is critical to safeguard correct gene expression profile during preimplantation development. Of interest, trimethylation of lysine 4 on histone 3 (H3K4me3) exhibits a unique and dynamic landscape with a potential species-specific feature. Here, we address how it is reprogrammed and its functional significance during oocyte maturation and early embryonic development in cows. Notably, the overall signal of H3K4me3 decreased sharply during embryonic genome activation (EGA). By using low input ChIP-seq technology, we find widespread broad H3K4me3 domains in oocytes and early cleaved embryos. The broad domains are gradually removed after fertilization, which is obviously seen during EGA. Meanwhile, H3K4me3 become enriched at promoter regions. Interestingly, the gene expression level displays a positive correlation with the relative H3K4me3 signal of their promoters when embryos reach 16-cell stage. Importantly, disruption of H3K4me3 demethylases KDM5A-5C increases H3K4me3 level, decreases the embryonic developmental rate and results in dysregulation of over a thousand genes. Meanwhile, KDM5 deficiency causes a re-destribution of H3K4me3 across genome. In particular, the positive correlation between promoter H3K4me3 enrichment and gene expression level disappear. Overall, we describe the genomic reprogramming of H3K4me3 in a greater resolution during bovine preimplantation development and propose that KDM5-mediated re-distribution of H3K4me3 plays an important role in modulating oocyte-to-embryonic transition.

## INTRODUCTION

The journey of a new life begins with the combination of oocyte and sperm in mammals. Upon fertilization, the highly differentiated gametes are magically reprogrammed to a totipotent embryo, when oocyte-to-embryo transition (OET) is completed (Rivera and Ross, 2013; Schultz et al., 2018; Xu and Xie, 2018). There are two major events occurring during OET: maternal mRNA clearance and embryonic genome activation (EGA)(Eckersley-Maslin et al., 2018; Schulz and Harrison, 2019). The time of EGA varies from species to species: mouse: 2-cell (Alberio, 2020), human: 4/8-cell (Niakan et al., 2012), pig: 4-cell (Whitworth et al., 2004), and cow: 8/16-cell stage (Graf et al., 2014; Misirlioglu et al., 2006).

Epigenetic modifications are dramatically reprogrammed and play an important role in orchestrate gene expression profiles during OET (Santos et al., 2002). As a hallmark of gene activation, the trimethylation of lysine 4 on histone H3 (H3K4me3) intensity is significantly reduced in accordance with EGA in various species, including mice, humans (Zhang et al., 2012), cows (Huang et al., 2015). However, the details of H3K4me3 reprogramming are just emerging thanks to the advent of low input epigenome methods, including ULI-NChIP-seq and CUT&RUN. Impressively, non-canonical broad H3K4me3 domains are enriched in gene bodies and intergenic regions during oogenesis and greatly removed shortly after fertilization in mice. Furthermore, the dysregulation of H3K4me3 removal appears detrimental for EGA and subsequent development (Dahl et al., 2016; Liu et al., 2016; Zhang et al., 2016a). Surprisingly, the genomic distribution of H3K4me3 in oocytes is quite different between human and mouse oocytes, suggesting species-specific role of H3K4me3 (Xia et al., 2019). Nevertheless, the landscape of H3K4me3 throughout bovine oocyte maturation and preimplantation embryogenesis as well as its functional role have yet to be determined.

In this study, we mapped genome-wide distribution of H3K4m3 in bovine oocytes and preimplantation embryos by taking advantage of low input ChIP-seq. Broad H3K4me3 domains were prevalent in bovine oocytes, and gradually erased after fertilization. Upon 16-cell stage, the broad H3K4me3 domains are completely removed while H3K4me3 is enriched at promoter regions. The positive correlation between the signal of promoter H3K4me3 and gene expression level becomes obvious from 16-cell stage. Besides, knocking down of KDM5A, KDM5B, and KDM5C in bovine embryos inhibited the erasure of broad H3K4me3 and blastocysts formation failure. In summary, we propose that timely removal of H3K4me3 is required for bovine early embryonic development.

## RESULTS

### Broad H3K4me3 domains are erased upon embryonic genome activation in cattle

To measure the global level of H3K4me3, we first collected bovine oocytes and preimplantation embryos and performed immunofluorescence (IF). Initially, relatively high level of H3K4me3 was observed in both GV and MII oocytes, but decreased significantly in 2-cell embryos. Astonishingly, H3K4me3 was barely seen at 16-cell stage (Fig 1a, 1b), when the major embryonic genome activation (EGA) occurs in cattle (Graf et al., 2014; Misirlioglu et al., 2006). The coincidence of H3K4me3 demethylation and EGA is also found in mouse, human and porcine embryos (Huang et al., 2015). Afterwards, H3K4me3 maintained low abundance in morulae and became brighter at blastocyst stage (Fig 1a, 1b). These results suggest H3K4me3 undergoes a substantial reprogramming after fertilization and, in particular, suffers a dramatical loss during EGA, which is conserved among species.

**Fig. 1.**
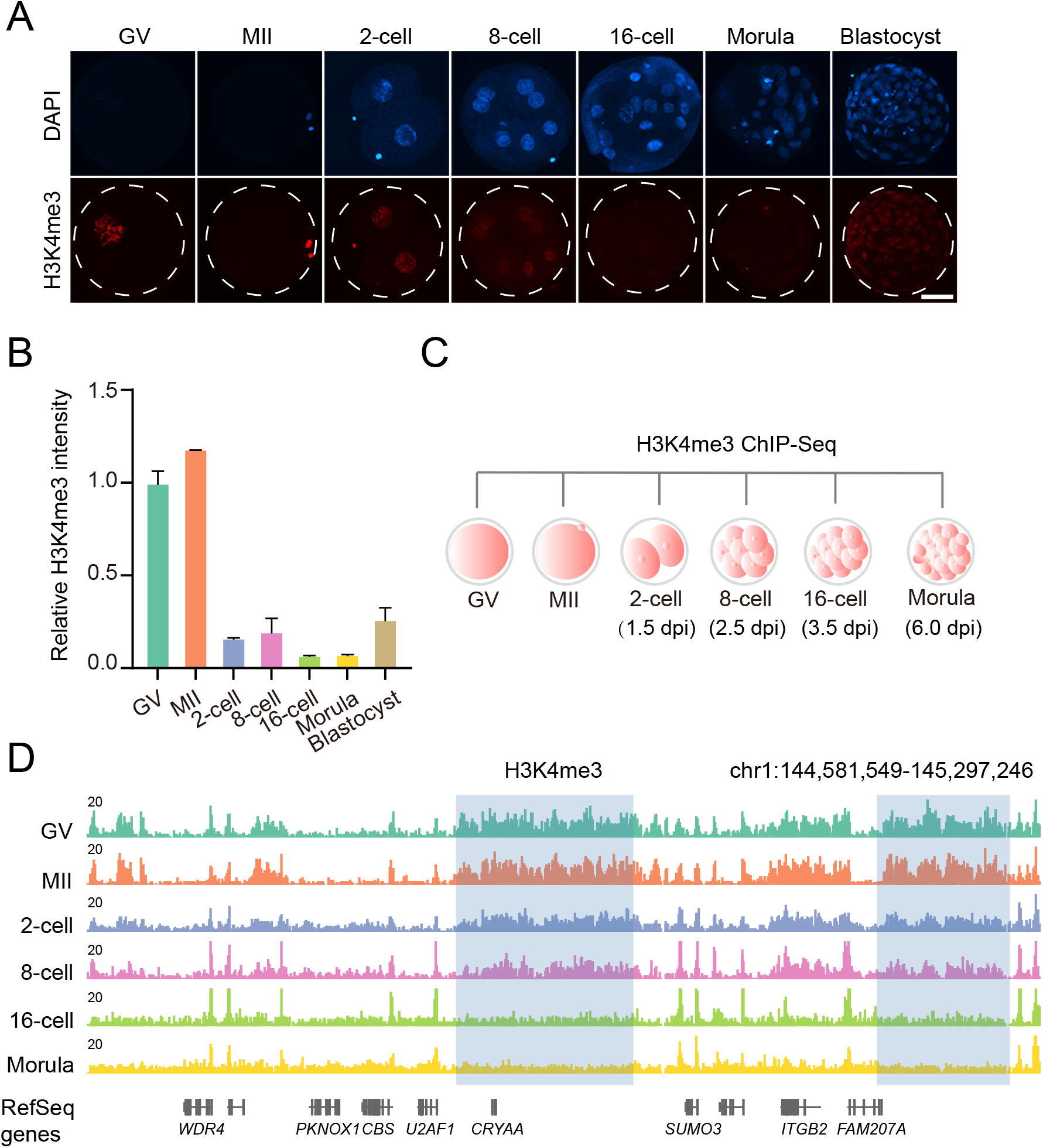
Dynamics of H3K4me3 during bovine early embryo development. **a,** Representative images for immunofluorescence staining of H3K4me3 in bovine oocytes and early embryos. Scale bar, 50 μm. **b,** H3K4me3 intensity relative to GV oocytes. Three replicates were conducted with at least 10 embryos in each stage, and data shown as mean ± SEM. **c,** Experimental scheme for H3K4me3 ULI-ChIP-seq. dpi, day post insemination. **d,** IGV browser snapshot for H3K4me3 distribution (two biological replicates were merged together).

In order to explore the genomic distribution of H3K4me3 in detail, we next performed ultra-low-input native chromatin immunoprecipitation (ULI-NChIP) in bovine oocytes and early embryos (Brind’Amour et al., 2015). To validate the protocol used here, we performed ULI-NChIP against H3K4me3 using 30 mouse morulae. Results showed that both biological replicates exhibited strong correlations, and recapitulated results from published data (R>0.80), indicating a robust reproducibility (Liu et al., 2016) (Fig S1a, S1b). Then, bovine oocytes and preimplantation embryos of 2-cell, 8-cell, 16-cell and morula stage were subjected to ULI-ChIP-seq (Fig 1c), and the data were reproducible between replicates (Fig S1c, S1d). Interestingly, we found broad/non-canonical H3K4me3 domains in both GV and MII oocytes, and the broad domains also existed in 2-cell and 8-cell embryos with slightly weaker signals but were almost completely removed in 16-cell embryos (Fig 1d). Besides, the broad H3K4me3 seemed to be more prevalent at gene body and intergenic regions (Fig 1d). Meanwhile, H3K4me3 signal at promoter region appeared to increase from 8-cell stage (Fig 1d). This removal of broad H3K4me3 domains is reminiscent of the similar H3K4me3 dynamics in mouse embryos (Dahl et al., 2016; Liu et al., 2016; Zhang et al., 2016a), but in sharp contrast to that in human embryos (Xia et al., 2019).

### Gene body or intergenic H3K4me3 is erased after fertilization and promoter H3K4me3 established during EGA

To test if broad H3K4me3 is truly prevalent at gene body and intergenic regions, we identified H3K4me3 peaks in oocytes and embryos of different stages, and classified them into gene body, intergenic and promoter peaks. Notably, the intergenic H3K4me3 distribution is very broad, which makes up ~ 13% of the genome in MII oocytes, however, the coverage decreased significantly after fertilization (Fig 2a). Moreover, the percentage of intergenic to total peaks reached its maximum value in MII oocytes and dropped constantly afterwards (Fig 2b). By scanning the genome with a 5 kb sliding window, we identified H3K4me3-lost and -gained regions in MII oocytes and embryos by comparing with the former stage. Genomic annotation analysis of H3K4me3-lost and -gained regions showed that more than 80% of H3K4me3-gained regions in MII oocytes were annotated as intergenic, while intergenic regions accounted for most of H3K4me3-lost rather than -gained regions in 2-cell, 8-cell, 16-cell and morula embryos (Fig S2a, S2b).

**Fig. 2.**
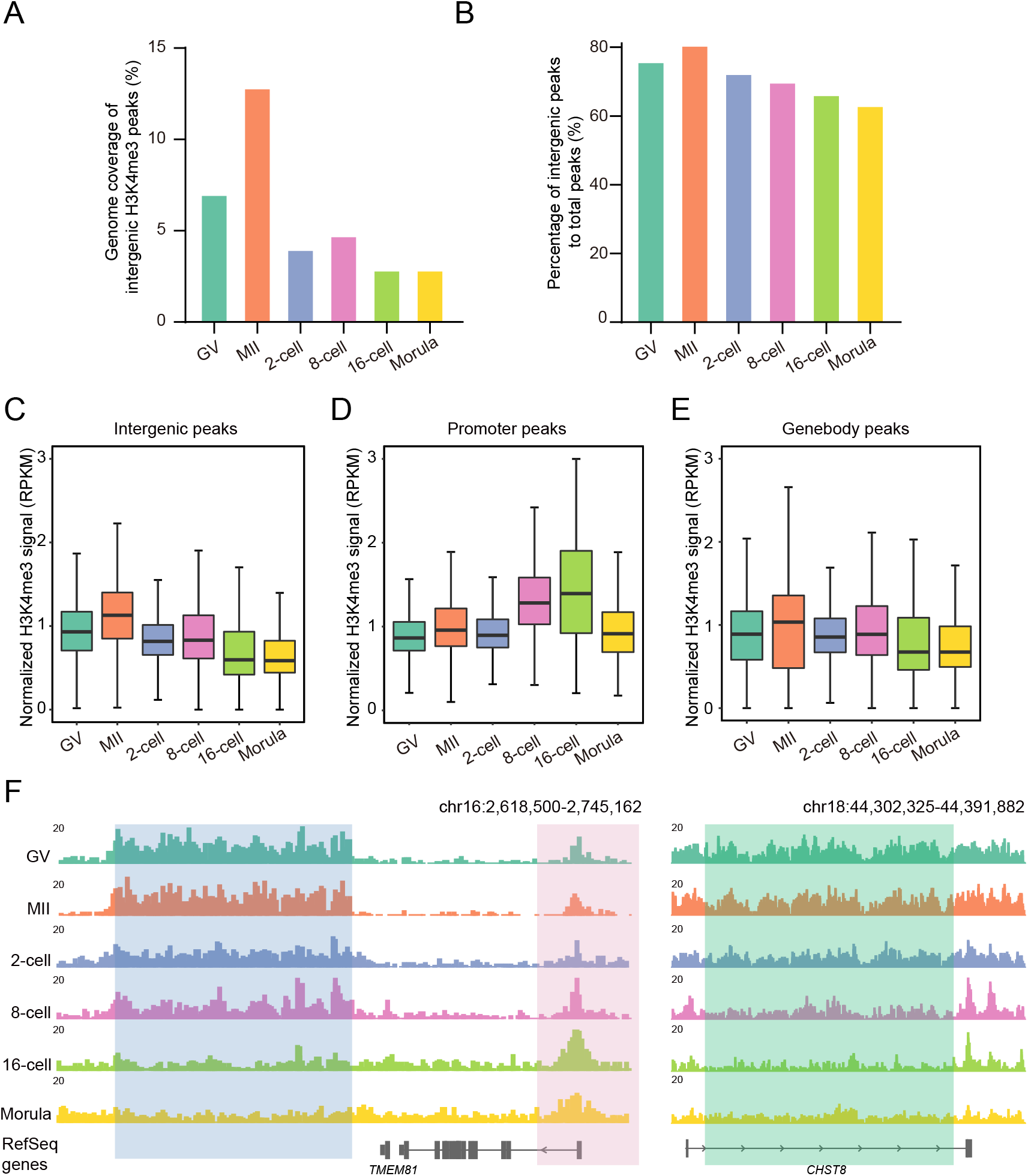
Distinct changes of H3K4me3 deposition at gene body, intergenic and promoter regions. **a,** Percentage of H3K4me3 peaks located in intergenic regions to the whole genome. **b,** Fraction of H3K4me3 peaks located in intergenic regions to total peaks. **c-e,** H3K4me3 signal of intergenic peaks (c), promoter peaks (d), and gene body peaks (e) in bovine oocytes and early embryos. **f,** IGV browser snapshot of H3K4me3 in different regions. The blue shadow refers to intergenic region, pink shadow refers to promoter region, and green shadow refers to gene body region.

To comprehensively investigate the genome-wide changes of H3K4me3, we then merged distal intergenic peaks together to calculate H3K4me3 enrichment at all potential intergenic regions. Distinctly, the signal of intergenic H3K4me3 peaks enhanced from GV to MII oocytes and became weaker at 2-cell and 8-cell stage, then continued to decrease in 16-cell embryos and morulae (Fig 2c, 2f). Similar with intergenic H3K4me3, gene body H3K4me3 also accumulated from GV to MII oocytes and was removed after fertilization (Fig 2e, 2f). Nevertheless, the signal of promoter H3K4me3 increased dramatically in 8-cell and 16-cell embryos (Fig 2d, 2f). These data collectively suggest broad H3K4me3 domains are established during oogenesis and greatly removed after fertilization. Meanwhile, the canonical H3K4me3 distribution at promoter regions are completed during EGA.

### Promoter H3K4me3 enrichment is positively associated with transcriptional activity upon EGA

H3K4me3 is well known as a hallmark of active promoters (Barski et al., 2007). Thus, we asked if H3K4me3 deposition is correlated with the abundance of transcripts in bovine early embryos. Based on the signal of promoter H3K4me3, the genes were divided into 2 clusters (Fig 3a). Genes in cluster 1 displayed increased enrichment of promoter H3K4me3 during 8/16-cell stage, while genes of cluster 2 lost promoter H3K4me3 at 16-cell stage (Fig 3a, 3b). Simultaneously, we quantified the average expression level of genes in the 2 clusters using published RNA-seq data (Graf et al., 2014). However, we found no correlation between promoter H3K4me3 enrichment and transcripts abundance in the 2 clusters (Fig 3c), suggesting increased enrichment of promoter H3K4me3 does not necessarily caused an increase in gene expression. Meanwhile, this result tell us again epigenetic regulation of gene expression is dependent on the developmental context, like different stages here.

**Fig. 3.**
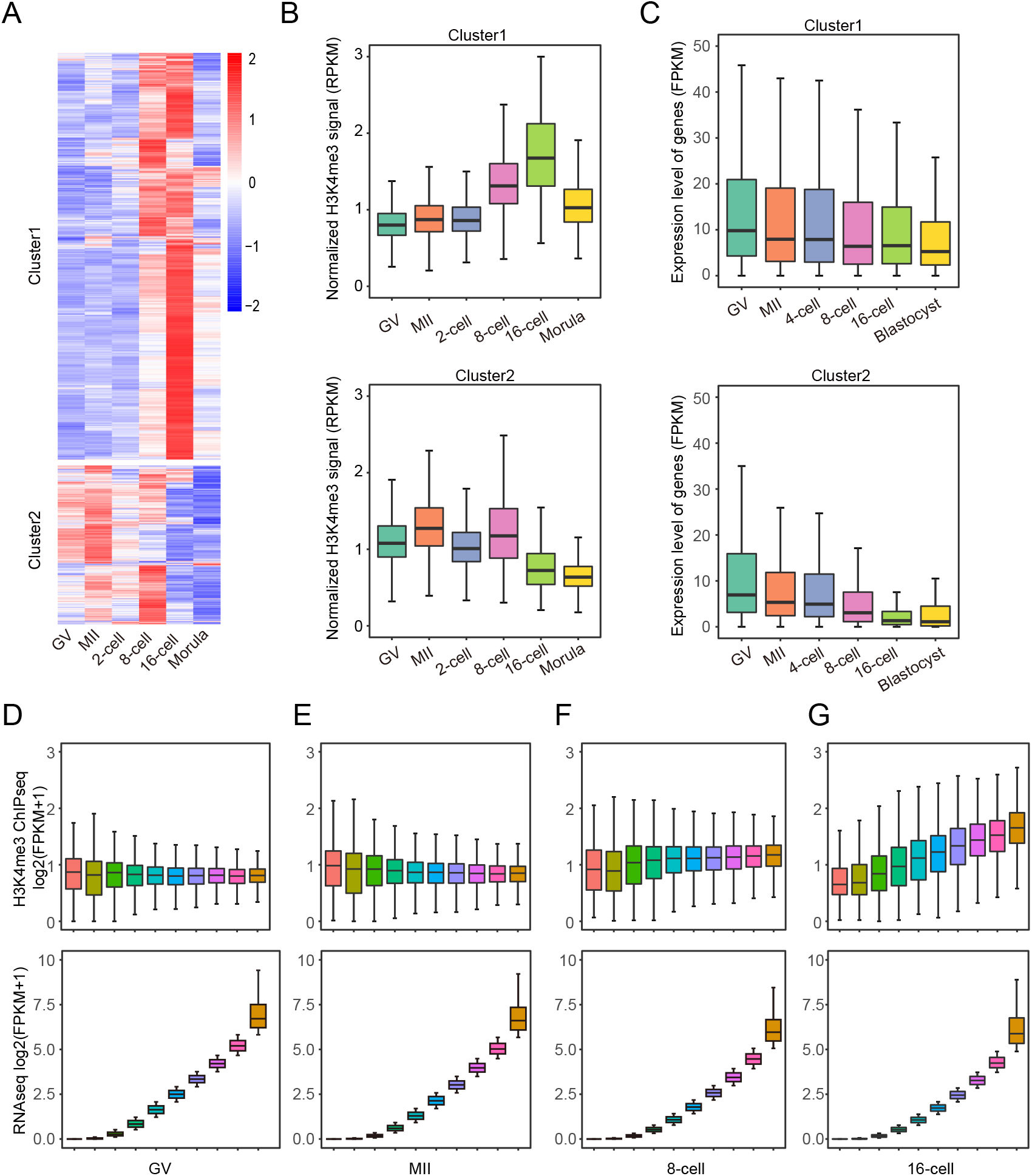
Promoter H3K4me3 could mark active genes especially at 16-cell stage. **a,** Heatmap showing z-score normalized RPKM of H3K4me3 in all promoter peaks. **b,** H3K4me3 signal of peaks in cluster 1 (top panel) and cluster 2 (bottom panel). **c,** Average gene expression level of genes in cluster 1 (top panel) and cluster 2 (bottom panel). The RNA-seq data was obtained from *Alexander Graf* et al. (Graf et al., 2014). **d-g,** H3K4me3 signal in promoter regions (top panel) and gene expression abundance (bottom panel) in the 10 equal sized groups.

We thus wondered if promoter H3K4me3 enrichment is correlated with gene expression within stages. We divided all genes into 10 groups by their expression abundance at each stage, and calculated the enrichment of promoter H3K4me3 in the corresponding group. In oocytes and early embryos prior to 8-cell stage, promoter H3K4me3 exhibits no or weak correlation with transcript abundance (Fig 3d-3f). Surprisingly, there was obvious positive correlation between the signal of promoter H3K4me3 and transcript abundance in 16-cell embryos (Fig 3g). Furthermore, the signal of H3K4me3 at gene body regions appeared negative correlation with gene expression in GV and MII oocytes (Fig S3), suggesting that H3K4me3 at gene body regions might be involved in transcriptional repression of oocytes.

### Interference with the removal of H3K4me3 by depleting KDM5A, KDM5B and KDM5C impairs bovine early embryonic developmental potential

Given the dynamic H3K4me3 reprogramming, we sought to explore the developmental and molecular consequence if the H3K4me3 erasure is blocked in bovine embryos. KDM5A, KDM5B and KDM5C are putative demethylases of H3K4me3 (Fig S4a). Interestingly, *KDM5B* and *KDM5C* showed a sharp increase in mRNA abundance during EGA, suggesting their functional requirement for the removal of H3K4me3 at 16-cell stage (Fig S4a). Our pilot study revealed KDM5A, KDM5B and KDM5C play a redundant role in H3K4me3 demethylation. Thus, we microinjected a cocktail of siRNA targeting all these three genes into zygotes for interfering the removal of H3K4me3 (Fig S4b). qPCR results indicated a robust knockdown efficiency (Fig 4a). As predicted, immunostaining data showed a significant increase of H3K4me3 in KDM5 co-knockdown (KD) embryos at both 8-cell and blastocyst stage (Fig 4b, 4c). Importantly, the proportion of 8/16-cell on day 3 post insemination was reduced in KD group (Fig 4d) and the blastocyst rate for KD group decreased significantly compared with NC group (Fig 4e). In all, the timely removal of H3K4me3 mediated by KDM5 is required for bovine early embryonic development.

**Fig. 4.**
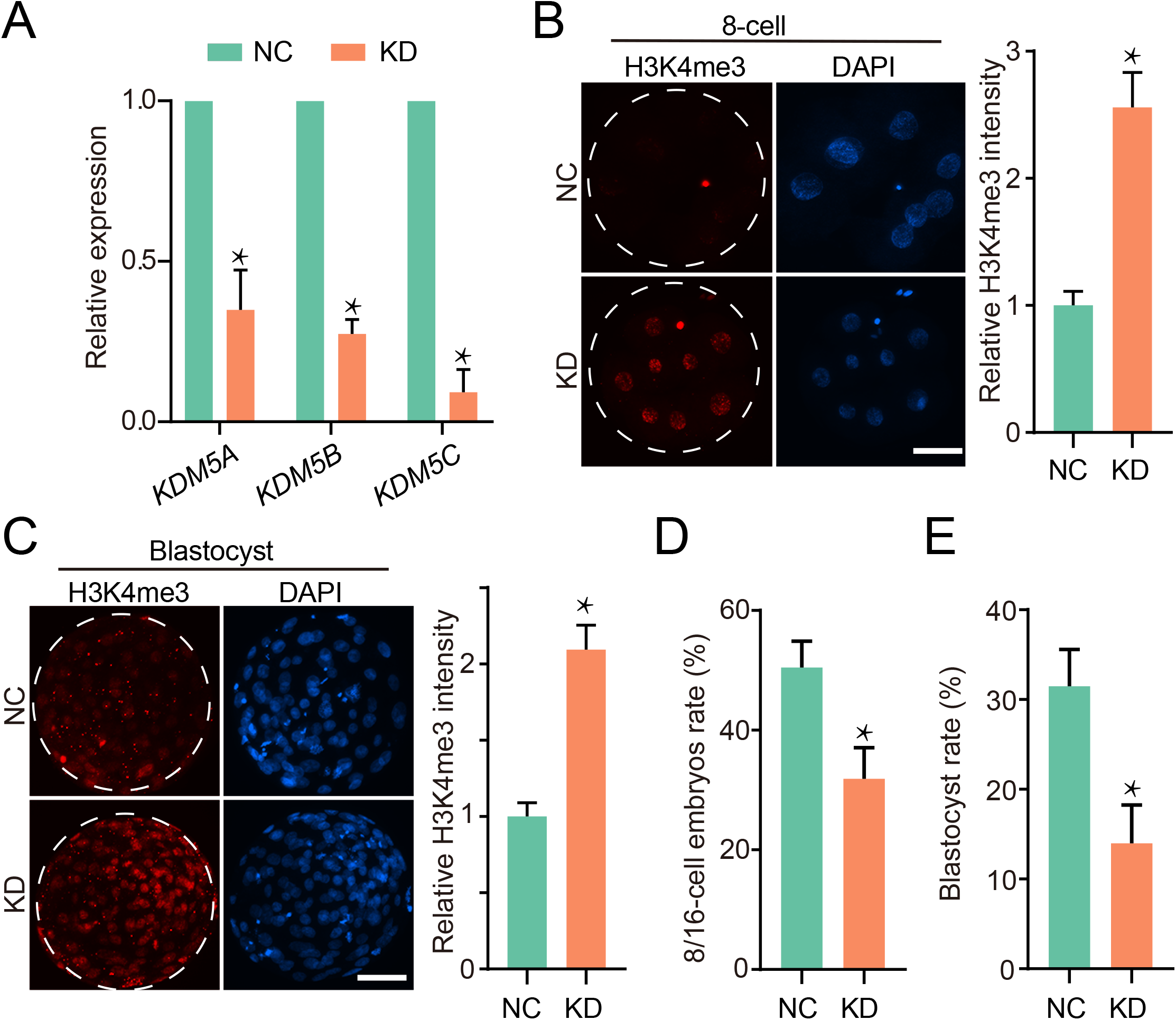
Depleting KDM5A, KDM5B and KDM5C prevented the removal of H3K4me3 and compromised blastocyst formation. **a,** Expression abundance of *KDM5A*, *KDM5B* and *KDM5C* for 8/16-cell embryos (3dpi) measured by qPCR. Three replicates were conducted with at least 15 embryos, and data was normalized to the abundance of endogenous control (*RPS18*) and shown as mean ± SEM. **b-c,** Immunofluorescence staining of H3K4me3 in 8-cell embryos (b) and blastocysts (c). Left, representative images, scale bar, 50 μm. Right, H3K4me3 intensity relative to negative control. Three replicates were conducted with at least 10 embryos in each stage, and data shown as mean ± SEM. **d-e,** Percentage of embryos developed to 8/16-cell (d) or blastocyst (e) monitored on 3 dpi or 7 dpi respectively. Three replicates were conducted with at least 20 embryos in each stage, and data shown as mean ± SEM.

### Blocked the removal of H3K4me3 dysregulated gene expression during EGA

In order to explore the effect of H3K4me3 removal on gene expression, we collected embryos from NC and KD groups at 8/16-cell stage to perform RNA-seq (Fig S4b). 1488 genes were differentially expressed in KD embryos, with 1310 up-regulated and 178 down-regulated (Fig 5a). We categorized the up-regulated and down-regulated genes into clusters based on their expression pattern in wild-type embryos. Among 178 down-regulated genes, 85 in cluster 1 are supposed to be highly expressed especially at 16-cell stage, and 50 in cluster 2 should be moderately expressed at 16-cell stage then highly expressed in blastocysts (Fig 5b). Only 24% (43/178) down-regulated genes (cluster 3) act as maternal genes, which are repressed during EGA (Fig 5b). For the 1310 up-regulated genes, 322 in cluster 5 are developmental genes which are prematurely expressed at 8/16-cell stage, and up to 826 in cluster 6 are maternal genes (Fig 5c). The remaining up-regulated genes (12%, 122/1310) can be classified as EGA genes (Fig 5c). The down-regulated genes were significantly enriched to several GO terms related to positive regulation of transcription activity as well as cell differentiation (Fig 5d). On the contrary, the up-regulated genes were enriched in transcriptional repressor complex and negative regulation of transcription. The results demonstrated that timely removal of H3K4me3 can regulate oocyte-to-embryonic transition via safeguarding the proper transcriptional activity.

**Fig. 5.**
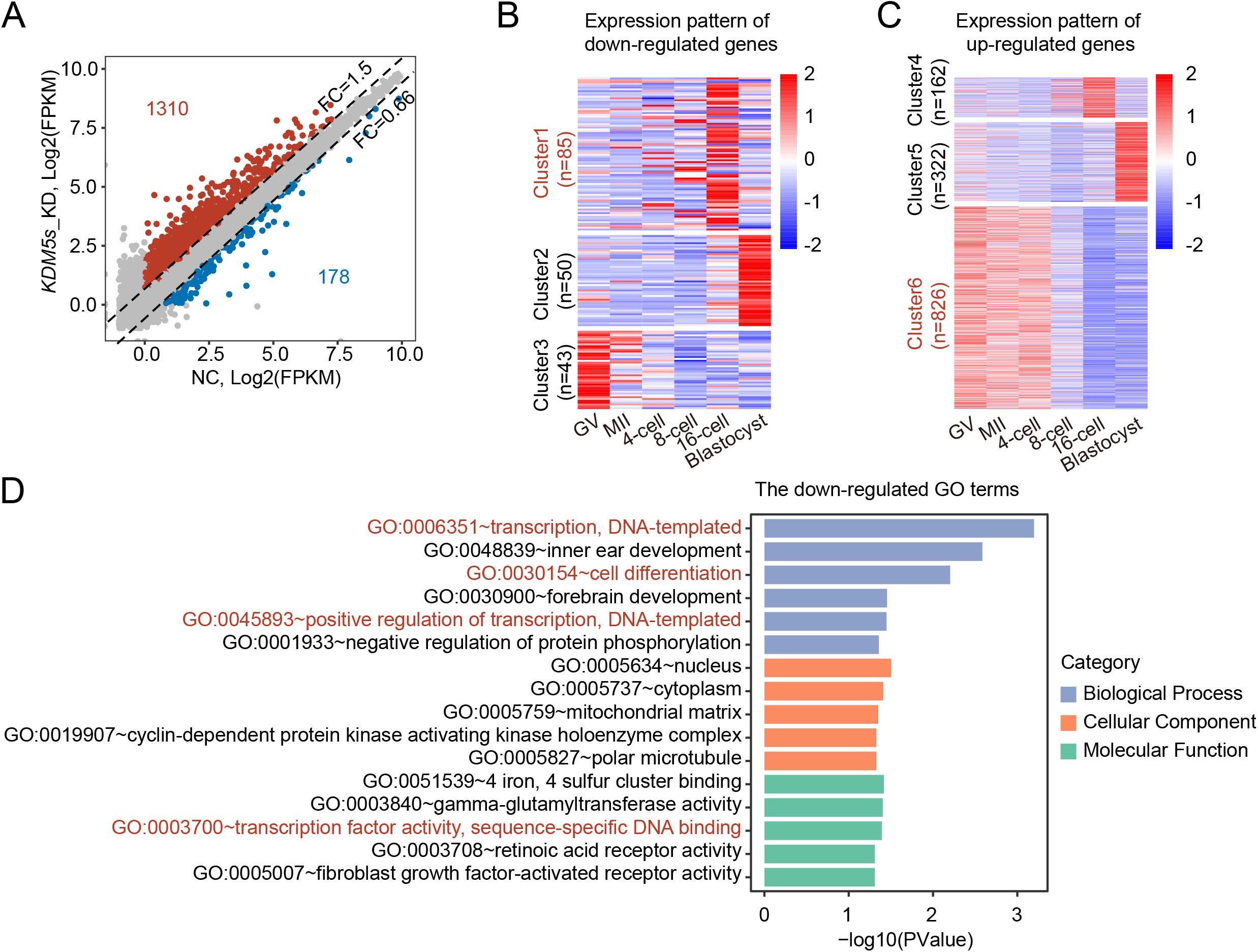
Disruption of H3K4me3 removal dysregulated gene expression in 8/16-cell embryos. **a,** Scatter plots showing gene expression level in NC and *KDM5s* KD group for bovine 8/16-cell embryos. Dash lines indicated the threshold of fold change (*KDM5s* KD/NC), and the up-regulated (*KDM5s* KD/NC>1.5) and down-regulated (NC/*KDM5s* KD>1.5) genes were colored with red and blue, respectively. Numbers of up- and down-regulated genes are also indicated in the figure. **b-c,** Heatmaps showing expression pattern of down- and up-regulated genes during wild-type bovine preimplantation development. The genes are classified into 6 clusters by *k*-means clustering. **d,** The enriched GO terms for down-regulated genes.

### Precise distribution of H3K4me3 is essential for transcription activity during EGA

We then tested that if the dysregulation of gene expression is attributed to the genomic re-distribution of H3K4me3 upon KDM5s knockdown. Thus, we performed H3K4me3 ULI-NChIP-seq at the same development stage with RNA-seq (Fig S4b). The two biological replicates for both groups exhibit high correlation (Fig S6a, S6b, S6c). By scanning the genome with a 5 kb sliding window, we identified H3K4me3-gained (Fig S6d) and -lost (Fig S6e) regions in KD embryos. H3K4me3-gained regions covered more genome than H3K4me3-lost regions (Fig 6a), which is in consistent with the IF results (Fig 4b). Further analysis revealed that more than 90% of the H3K4me3-gained regions were gene-body or intergenic regions (Fig 6b, 6c), indicating the removal of broad H3K4me3 in these regions was largely impaired. Indeed, gene body or intergenic H3K4me3 peaks covered more genome in the KD embryos (Fig S6f).

**Fig. 6.**
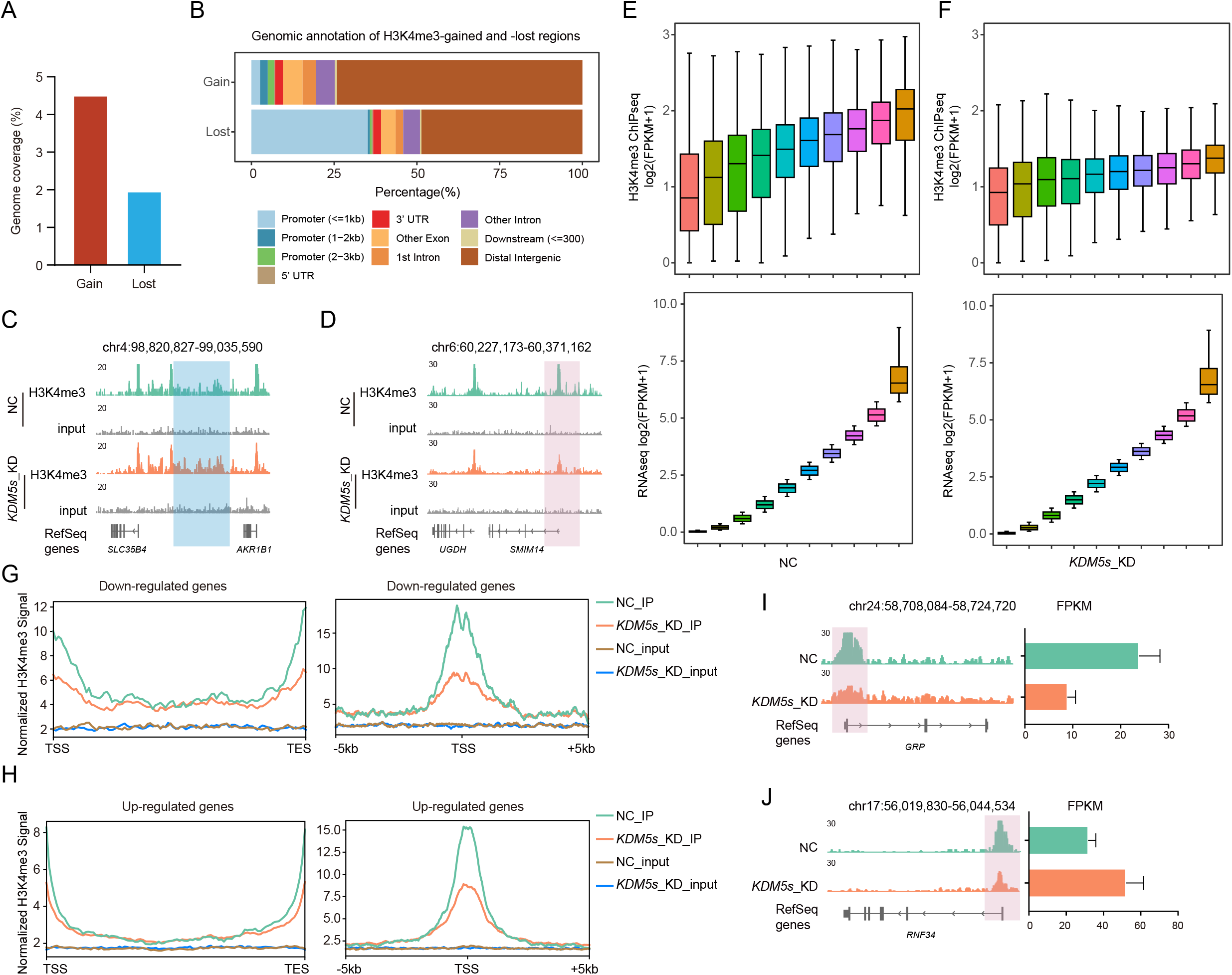
Depleting KDM5A, KDM5B and KDM5C changed the genomic distribution of H3K4me3. **a,** Genomic coverage for H3K4me3-gained and -lost regions. Identification of H3K4me3-gained and -lost regions are described in the Methods section. **b,** Genomic annotation for H3K4me3-gained and -lost regions. **c-d,** IGV snapshot for H3K4me3-gained (c) and -lost regions (d). **e-f,** Gene expression abundance (bottom panel) and H3K4me3 signal at promoter regions of the corresponding genes (top panel) in NC (e) and *KDM5s* KD group. **g-h,** Left panel showed profiles of H3K4me3 signal in gene body regions of all down-regulated (g, NC/*KDM5s* KD >2) and up-regulated (h, *KDM5s* KD/NC >2) genes. The right panel shows profiles of H3K4me3 signal at promoters of all down-regulated genes (g) and up-regulated genes (h). H3K4me3 signal was normalized to RPKM by DeepTools. **i-j,** IGV browser snapshots for down- (i) and up-regulated gene (j).

Genome annotation of H3K4me3-gained regions showed ~40% of them were located in promoter regions (Fig 6b, 6d), reflecting the inadequate establishment of promoter H3K4me3. In addition, the average signal of H3K4me3 in promoters of all genes decreased significantly in the KD embryos. These results demonstrated that KDM5 is not only required for the timely removal of broad H3K4me3 in gene body and intergenic regions but the establishment of promoter H3K4me3.

Since promoter H3K4me3 enrichment is gradually correlated with gene expression especially after the removal of broad H3K4m3 during EGA (Fig 3d-3g), we then tested if it is still the same case upon KDM5 KD. All genes expressed in NC or KD embryos were divided into 10 groups equal in number according to their abundance, and the enrichment of promoter H3K4me3 was quantified in each group respectively. The result showed that correlation between gene expression level and promoter H3K4me3 signal became weaker in KD group.

## DISSCUSSION

Although substantial reprogramming of histone modifications is critical to mammalian early embryonic development, the investigation of the underlying mechanisms has been hindered by the lack of chromatin analysis approach in sparse samples. Recently, the advent of low input epigenome approach, including ChIP-seq and Cut & Run, has allowed us to look into the reprogramming of histone modifications in great details. In the present study, we describe the dynamic distribution of H3K4me3 across genome and address its functional importance during oocyte maturation and early embryonic development in cattle.

Global H3K4me3 signal decreased during mouse, porcine and human EGA (Dahl et al., 2016; Huang et al., 2015; Zhang et al., 2012). Our work found a similar change in bovine embryos, indicating the overall removal of H3K4me3 is conserved among species. Utilizing ULI-NChIP-seq technology, we profiled the genome-wide mapping of H3K4me3 in bovine oocytes and preimplantation embryos, and observed widespread broad H3K4me3 regions in both oocytes and embryos before EGA, which is similar to mouse but in contrast to those in human (Dahl et al., 2016; Xia et al., 2019; Zhang et al., 2016a). Similar with mouse model, we found prevalent distribution of broad H3K4me3 at gene body and intergenic regions in oocytes, which was removed sharply during EGA (Fig 7). Importantly, in contrast with the overall loss of H3K4me3 detected by IF, ChIP-seq analysis revealed enhanced H3K4me3 at promoter regions during EGA (Fig 7). This reminds that when studying dynamics of epigenetic modification, the combination of IF and ChIP-seq is very important to obtain accurate and comprehensive information.

H3K4me3 has been considered as a hallmark of actively transcribed genes. However, its relationship with transcription activity varies in different research models (Howe et al., 2017). Meanwhile, the broad/non-canonical H3K4me3 at non-promoter regions seen in oocytes and early cleaved embryos further complicates the functional role of H3K4me3 on transcription. Here, the H3K4me3 signal at promoter regions showed an obviously positive correlation with gene expression only when broad H3K4me3 domains was removed upon EGA. It is possible that broad H3K4me3 is required for protecting genome from premature transcription as seen in mouse oocytes. Subsequently, the removal of these broad domains and enrichment of promoter/canonical H3K4me3 is required for setting up transcriptional program during EGA. However, H3K4me3 per se seems to play a limited role in transcription activation as KDM5-medidated H3K4me3 redistribution only results in dysregulation of limited number of genes. This result is consistent with our recent finding that H3K4me3 level is not a reliable marker to distinguish active and repressive genes in mouse early embryos (Dang et al., 2021). The crosstalk between H3K4me3 and other histone modifications such as histone acetylation H3K27ac may be a potential mechanism that recruits other chromatin factors to contribute to the eventual transcription activation.

In summary, we characterize the genome-wide distribution of H3K4me3 in a greater resolution using the cutting-edge ChIP-seq approach and reveal the dynamic changes of H3K4me3 and its association with gene expression during early embryonic development in bovine model. Importantly, we show the functional requirement of KDM5-mediated H3K4me3 demethylation in bovine early embryonic development.

## MATERIALS AND METHODS

### *In vitro* production of bovine embryos

Bovine oocyte in vitro maturation (IVM), in vitro fertilization (IVF) and embryo in vitro culture (IVC) were performed according to procedures as published previously with slight modifications (Zhang et al., 2015; Zhang et al., 2016b). Briefly, we collected bovine ovaries from a local slaughterhouse. Cumulus oocyte complexes (COCs) containing more than three layers of cumulus cells were retrieved from 3–8-mm follicles at the surface of bovine ovaries. COCs were cultured in Medium-199 (M4530) supplemented with 10% FBS (Gibco-BRL), 1 IU/ml FSH (Sansheng Biological Technology), 1 mM Sodium Pyruvate (Thermo Fisher Scientific), 2.5 mM GlutaMAX™ (Thermo Fisher Scientific), and 10 μg/ml Gentamicin at 38.5°C under 5% CO_2_ in humidified air for 22-24 h. Upon maturation, COCs (100 COCs/ well in 4-well plates) were co-incubated with spermatozoa (1×10^6^) that purified from frozen-thawed semen with a Percoll gradient. IVF was performed at 38.5°C under 5% CO_2_ for 9-12 hrs. Cumulus cells were discarded from the oocytes by pipetting up and down with 1 mg/ml hyaluronidase. Embryos were cultured in BO-IVC medium (IVF bioscience) for 8 days or collected until use.

### Immunofluorescence (IF)

Samples were briefly washed with 0.1% polyvinylpyrrolidone/PBS three times, fixed with 4% paraformaldehyde/PBS for 30 minutes, and permeabilized with 0.5% Triton X-100/PBS for 30 minutes at room temperature. Embryos were blocked for 1 h in PBS containing 10% FBS and 0.1% Triton X-100. Then, samples were incubated with primary antibodies in blocking buffer under 4°C overnight and secondary antibodies for 2 h. Finally, samples were treated with DAPI (Life Technologies) for 30 minutes, and then mounted onto slides. Images were captured with a 40× objective using an inverted epi-fluorescent microscope (Nikon). ImageJ (https://imagej.nih.gov/ij/index.html) was used to visualize images, count cell numbers, and measure signal intensity. For certain experiments, the signal intensity was determined and the background subtracted to analyze the absolute intensity.

### RNA interference in early embryos

RNA interference in bovine early embryos was performed as published (Zhang et al., 2015; Zhang et al., 2016b). siRNA oligo sequences were synthesized by GenePharma (Shanghai, China). Basic Local Alignment Search Tool (BLAST; https://blast.ncbi.nlm.nih.gov/Blast.cgi) was used to rule out potential non-specific targeting. A cocktail of three siRNAs was used per gene of interest to maximize the interference efficiency. In control groups, embryos were microinjected with non-specific siRNAs at the same concentration. To validate the efficacy of siRNAs used, 8/16-cell stage embryos were collected for qPCR analysis. Immunofluorescence was also performed to validate knockdown effects at the protein level.

### RNA isolation, reverse transcription, and quantitative PCR

Total RNA from 8/16-cell embryos (10-15/group/replicate) was isolated using the Arcturus PicoPure RNA Isolation Kit (Applied Biosystems). Purified RNA was then subject to reverse transcription with SuperScript II Reverse Transcriptase (Invitrogen) and quantitative PCR (qPCR) conducted on a StepOne system (Applied Biosystems) using FastStart Universal SYBR Green Master (Roche). Quantification was normalized to the endogenous control (*RPS18*). One embryo equivalent of cDNA was used for each qPCR reaction with a minimum of three replicates for the data shown.

### ULI-NChIP-seq

GV and MII oocytes were collected before and after maturation. Bovine embryos were harvested at the following time: 1 day post insemination (dpi) for 2-cell, 3 dpi for 8-cell, 4 dpi for 16-cell, and 5 dpi for morulae. For KDM5 knockdown experiment, embryos were collected at 8/16-cell stage (3 dpi). Oocytes and embryos were incubated in 0.5% pronase E (Sigma) for 3-5 minutes to remove zona pellucida, and washed 3 times in 0.5% bovine serum albumin (Millpore) in DPBS (Gibco) before flash freezing in liquid nitrogen. ULI-NChIP was performed according to the published step-by-step procedure (Brind’Amour et al., 2015). 1 μg H3K4me3 (CST 9751) was used for each immunoprecipitation reaction. ULI-NChIP libraries were generated using the NEB Ultra DNA Library Prep Kit (E7645). The libraries were then sent to Novogene Co., Ltd for a paired-end 150 bp sequencing on NovaSeq (Illumina) platform.

### RNA-seq

Embryos were harvested at 8/16-cell stage (3 dpi). Before RNA isolation, equal amount of *GFP* and *RFP* mRNA were added to each sample as a spike-in control. Total RNA was extracted using Arcturus PicoPure RNA Isolation Kit (Applied Biosystems). Then mRNAs were separated with oligo(dT)25 beads. Sequencing libraries were built with NEB Next Ultra RNA Library Prep Kit for Illumina (E7530). Briefly, mRNAs were fragmented followed by reverse transcription and then underwent end repair, poly(A)-tailing, adaptor ligation, and PCR amplification for 12–15 cycles to prepare sequencing libraries. Libraries were sent to Novogene Co., Ltd for a paired-end 150 bp sequencing on NovaSeq (Illumina) platform.

### ULI-NChIP-seq data processing

The raw sequencing reads were trimmed with Trimmomatic (version 0.39) (Bolger et al., 2014) to remove residual adapter sequences and low-quality bases. The clean reads were aligned to Bos Taurus UMD3.1.1 using Bowtie2 (version 2.3.5) (Langmead and Salzberg, 2012) using default parameters. Mapped reads with low quality were removed by SAMtools (version 1.7)(Li et al., 2009), and PCR duplicates discarded with Picard (version 2.23). To visualize H3K4me3 signal in IGV genome browser, the genome was binned into 50 bp windows and RPKM for each window calculated using bamCoverage from DeepTools (Ramirez et al., 2014).

### RNA-seq data processing

The raw sequencing reads were trimmed with Trimmomatic (version 0.39) to generate clean data, and clean reads were then mapped to Bos Taurus UMD3.1.1 with Hisat2 (version 2.1.0)(Kim et al., 2015). The raw counts were calculated with featureCounts (version 1.6.3)(Liao et al., 2014) and FPKM for each sample calculated with Cufflinks (version 2.2.1)(Trapnell et al., 2012). The differential expressed genes were identified when the fold change is greater than 1.5 (Dahl et al., 2016; Matoba et al., 2014). For clustering the differential expressed genes, RNA-seq data of bovine oocytes and embryos was obtained from Alexander Graf et al (Graf et al., 2014). The FPKM matrixes were generated with Cufflinks as described above. Then the non-zero FPKM of genes was used for *k*-means clustering, which was performed with R based on Euclidean distance. GO enrichment analysis of differentially expressed genes was performed with the Database for Annotation, Visualization and Integrated Discovery (DAVID)(Huang da et al., 2009a, b).

### Reproducibility of ULI-ChIP-seq data

The genome was binned into 5000bp windows using makewindows utility of Bedtools (Quinlan and Hall, 2010), and the reads counts of each window calculated and normalized to RPKM. Then the RPKM value for each replicate was used to calculate Pearson correlation coefficient and plotting by R. The correlation between our data and the published data (Liu et al., 2016) was measured in the same way as above.

### Analysis of H3K4me3 peaks at genebody, intergenic and promoter regions

Since the number of peaks are affected by sequencing depth and equal numbers of ChIP and input reads result in best performance of peak callers (Chen et al., 2012; Jung et al., 2014), we merged alignments of biological replicates and randomly subsampled them to equivalent depth using SAMtools before peak calling. H3K4me3 peaks were called with MACS2 (Zhang et al., 2008) using the following parameters: -B -p 1e-5 --nomodel --broad --extsize 73. Those peaks were then annotated using ChIPseeker (Yu et al., 2015), and peaks annotated as UTR, intron or exon regions were collectively considered as genebody H3K4me3 peaks. The annotated distal intergenic regions were intergenic peaks, and peaks within 3kb of TSS were promoter peaks.

### The correlation between gene expression level and promoter H3K4me3

Genes in each stage were divided into 10 equal groups according to expression levels, then the H3K4me3 distribution at promoter regions in corresponding group was normalized to RPKM.

### Identification of H3K4me3-gained or -lost regions

To compare H3K4me3 distribution between groups, bovine genome was scanned using a sliding window of 5 kb and step size of 1 kb. H3K4me3 signal for each window was calculated and normalized to RPKM. Next, RPKM of H3K4me3 was compared in parallel between groups. The H3K4me3-gained or -lost regions were identified with following threshold: log2 (fold change) > 1.5 for gained regions or < −1.5 for lost regions, and sum of RPKM in groups >1. Then the selected regions were merged with BEDTools if they were overlapped, and the genome coverage of merged regions was calculated by genomecov mode of BEDTools.

### Dataset statement

Bovine RNA-seq data for wild-type embryos are obtained from the published data (GSE52415). The sequencing data underlying this article have been deposited to GEO with the accession number GSE189102.

### Statistical analyses

Three biological replicates were conducted unless stated. Two-tailed unpaired Student’s t test were performed to compare differences between groups. Immunofluorescent intensities were measured with ImageJ. Figures were generated by using GraphPad Prism 8 (GraphPad Software). Value of P<0.05 denotes statistical significance. Results are shown as means ± SEM.

## Acknowledgments

We thank all members of the K. Zhang laboratories for their helpful discussions. This work was funded by National Natural Science Foundation of China (No. 31872348, No. 31672416, and No. 32072731 to K.Z.; No.31941007 to L.L. and S.W.), Zhejiang Provincial Natural Science Foundation (LZ21C170001 to K.Z.) and China Postdoctoral Science Foundation (No. 2020M671742 to L.L.).

---

**Fig. 7 Model illustrating dynamics of H3K4me3 distribution and the role of KDM5s.**

During GV to MII stage, H3K4me3 deposited at gene body or intergenic regions increased moderately. While after fertilization, the deposition of H3K4me3 in gene body or intergenic regions was greatly removed and almost disappear at 16-cell stage by KDM5A/B/C. Notably, H3K4me3 signal in promoter regions was established along with the removal of distal H3K4me3, and KDM5s contributed to the establishment of promoter H3K4me3 in an indirect way. Besides, although H3K4me3 has distinct changes during EGA, H3K4me3 alone has only a very limited effect on EGA, and it may need to cooperate with other histone modifications (eg, H3K27ac) to regulate the transcriptional activity during this process.

